# Endocytic motif on HIV-1 Env regulates cleavage status and antibody neutralization of cell-free and cell-to-cell infection

**DOI:** 10.1101/2025.09.15.676328

**Authors:** Dania M. Figueroa Acosta, Hongru Li, Benjamin K. Chen

## Abstract

Antibodies inhibit human immunodeficiency virus type 1 (HIV-1) infection by targeting the envelope glycoprotein (Env). Env cleavage is a key determinant of antibody binding, as cleavage reduces Env flexibility and alters glycosylation. Given the greater abundance of uncleaved Env on the cell surface compared to virions, we investigated whether Env endocytosis, initiated through a membrane-proximal tyrosine motif in its cytoplasmic tail, regulates the abundance of cleaved Env at the cell surface. We hypothesize that such a shift would alter the cleavage and glycosylation profile of cell surface Env, affecting the sensitivity of cell-to-cell infection to neutralizing antibodies. To address this, we generated an endocytic mutant (ASPI-Env) and compared its antigenic properties to a cleavage-site mutant (SEKS-Env). Immunoprecipitation and ratiometric antibody binding studies of cell surface Env demonstrated that the ASPI mutation increases the amount of uncleaved Env on the cell surface. Consequently, ASPI-Env in cell-free infection was more sensitive to NAbs recognizing uncleaved Env. Notably, only b12, which can engage both cleaved and uncleaved Env but has a higher affinity for uncleaved Env, showed increased inhibition of ASPI-Env during cell-to-cell infection. This mirrors the enhanced neutralization of SEKS-Env during the cell-to-cell transfer assay and supports a model in which uncleaved Env participates in CD4-dependent virion transfer. Finally, the ASPI mutation altered lectin binding and differentially affected cell-to-cell and cell-free infections. Together, these findings indicate that Env endocytosis modulates the abundance of cleaved Env at the cell surface, thereby influencing antibody neutralization and lectin recognition in distinct modes of HIV-1 transmission.

**IMPORTANCE:** We find that Env endocytosis modulates the antigenicity of Env on both the cell surface and virions. Blocking internalization increased uncleaved Env on the cell surface, thereby reshaping HIV-1 neutralization due to enhanced binding of antibodies that preferentially recognize uncleaved Env. We further show that uncleaved Env on the cell surface can initiate CD4-dependent transfer of virions across virological synapses. These findings demonstrate that Env endocytosis shapes cleavage-associated antigenicity in a manner that can differentially impact the recognition and neutralization of cells and viruses. The impact of uncleaved Env on neutralization indicates that a broader spectrum of cleaved and uncleaved Env conformations may be relevant when designing vaccines and cure strategies that must contend with a diverse antigenic landscape.

## INTRODUCTION

The human immunodeficiency virus infects CD4_+_ T cells through both cell-free virus and direct cell-to-cell routes (1). Cell-free infection occurs when viral particles released from an infected cell diffuse and infect target cells. In contrast, cell-to-cell infection is initiated when Env, expressed on the surface of an infected cell, binds to the CD4 receptor of a neighboring cell, thus triggering the formation of a virological synapse (VS) (2–4). This facilitates a local high multiplicity of infection as multiple virions are transferred and either fuse at the plasma membrane or are internalized via endosomes (4–8). Although the viral envelope glycoprotein (Env) mediates both mechanisms of infection, *in vitro* studies have shown that cell-to-cell infection is more resistant to inhibition by broadly neutralizing antibodies (bNAbs) than cell-free infection (3, 9–11). This resistance persists even at high bNAb concentrations, resulting in incomplete neutralization of cell-to-cell infection, in contrast to the complete neutralization of cell-free virus (12). The reduced sensitivity of cell-to-cell infection to bNAbs is thought to reflect differences in the antigenic state of Env during the two modes of infection (9, 13).

The application of single-molecule Förster resonance energy transfer (FRET) to understand conformational dynamics of the Env glycoprotein reveals that Env is a conformationally dynamic protein that spontaneously transitions between three distinct prefusion conformations (14). These states, a closed, stable ground-state, an asymmetric intermediate, and an open trimer conformation, are the same conformations Env adopts upon CD4 binding (14, 15). Independent of CD4 binding, the cleavage status of Env influences its conformational state: cleavage decreases Env’s conformational flexibility, causing it to predominantly adopt the closed, ground-state configuration (16, 17). In contrast, uncleaved Env samples all three conformations more freely (16, 17). Consequently, cleavage can alter epitope exposure and antibody binding (18–21). Cleavage has also been implicated in modulating Env’s glycan composition by limiting access to glycan processing enzymes in the Golgi (22). Given that glycans account for roughly 50% of Env’s mass and since many bNAbs are glycan-dependent (23–27), Env’s cleavage state can significantly affect bNAb neutralization potency.

Env cleavage is variable, resulting in a mixture of cleaved and uncleaved Env being expressed on the cell surface (28, 29). However, viral particles are highly enriched for cleaved Env (29). The mechanism for such selective incorporation of cleaved Env into virions remains largely undefined, though Env’s cytoplasmic tail (CT) has been implicated in this process as truncations of the CT reduce infectivity (30, 31). The CT has many motifs, including a membrane-proximal tyrosine-based motif (YXXΦ), which is responsible for clathrin-mediated endocytosis of Env (32–34). Previously, Li et al. observed that mutation of the endocytic motif altered the neutralization sensitivity of cell-free and cell-to-cell infection in an antibody-dependent manner (12). Similarly, Chen et al. demonstrated that truncations in Env’s CT altered Env’s ectodomain, revealing epitopes for non-neutralizing antibodies (35). Data from analogous mutants in SIV have documented attenuated replication and delayed depletion of mucosal T cells when the endocytic motif was mutated (36, 37). These findings suggest that Env’s endocytic motif shapes the epitope exposure of Env engaged in HIV-1 infection and contributes to HIV-1 immune evasion.

In this study, we explored how active endocytosis contributes to the antigenic state of cell surface Env by modulating the abundance of cleaved Env and its glycosylation pattern, thus promoting differences in the potency and efficacy of bNAbs in cell-free and cell-to-cell infection. To test the hypothesis that sorting affects immune recognition of Env, we generated an endocytic-deficient Env (ASPI-Env) from the T/F clade B HIV-1 QH0692. To determine whether its antigenic profile resembled a more open conformation representative of uncleaved Env, we also produced a furin-resistant mutant Env (SEKS-Env). We assessed the cleavage and glycosylation states of the wildtype (WT) and mutant Envs using a panel of monoclonal antibodies and lectins. Immunoprecipitation of cell-surface Env and a ratiometric antibody binding assay revealed that ASPI-Env expressed on the cell surface mimicked the antigenic profile of SEKS-Env-expressing cells. To assess the functional consequences of these conformational changes, we performed neutralization assays for both cell-free and cell-to-cell infections. Finally, to further understand the influence of uncleaved Env conformation on cell-to-cell infection, we also quantified the transfer of fluorescent viral particles. Our study reveals that endocytosis contributes to the antigenic heterogeneity of Env involved in HIV-1 infection and affects its recognition by neutralizing antibodies.

## MATERIALS AND METHODS

### HIV-1 Env variants

The transmitted/founder (T/F) HIV-1 QH0692 Env (ARP Cat #11227, Drs. David Montefiori and Beatrice Hahn) was cloned into the mCherry fluorescent protein-expressing pNL4-3-based molecular clone NL-CI_NL4-3_ (38) and Gag-iCherry fusion protein expressing clone, HIV-1 Gag-iCherry as previously described (39). To generate an endocytic mutant of QH0692 Env, we introduced a mutation at the main endocytic motif site residue Y724A, from YSPI to ASPI by PCR amplification using CloneAMP (Takarabio, Cat. 639298). We also generated a cleavage-defective Env mutant by mutating the primary cleavage site residues R506S and R509S, from REKR to SEKS. Following digestion of the pNL4-3 plasmid with EcoRI-HF and MluI-HF, an In-Fusion reaction was performed with the In-Fusion HD Cloning Kit (Clontech Labs, Cat. 3P639648). HIV-1 Gag-iCherry was digested with EcoRI-HF and XhoI-HF, followed by Gibson assembly (New England BioLabs, Cat. E5510S). The mutants were confirmed by Sanger sequencing across all PCR-amplified sequences.

### Cell culture

293T cells were obtained from the American Type Culture Collection (ATCC). RevCEM-D4 (ARP-13437) and Jurkat E6 (ARP-177) cells were obtained from the NIH Reagent Program from Dr. Alex Sigal and Dr. Arthur Weiss, respectively. Primary CD4_+_ T cells were isolated from human peripheral blood through the New York Blood Center. All cells were cultured as previously described in Durham et al. and Li et al. (12, 40). Primary CD4_+_ T cells were enriched from peripheral blood mononuclear cells (PBMCs) using CD4_+_ T cell Isolation Kit according to the manufacturer’s instructions (Cat. 130096533, Miltenyi Biotec) and activated by supplementing media with 4µg/mL of Phytohemagglutinin-L (SigmaAldrich, Cat. 11249738001) and 50IU/mL of IL-2 (Miltenyi Biotec, Cat. 130097746). Cell-free virions were produced by transfecting 293T cells, as described in Li et al. (12). After 48 hours, cells were harvested for western blotting and supernatant for virus.

### Virus Production and immunoprecipitation of cell surface Env

293T cells were transfected using PolyJet In Vitro DNA Transfection Reagent (SignaGen Laboratories, Cat. SL1006885X1ML). After 48 hours, cells were collected by centrifugation at 500xg, while the virus was spun at 1000xg to remove debris and filtered through a 0.45μM filter. Immunoprecipitation was adapted from Zou et al.’s protocol to isolate cell-surface Env (28). Briefly, cell samples were incubated with 10µg/mL monoclonal antibodies PGT151, b12, 35O22, 17b and 2G12 for 1 hour at 4°C. The following antibodies were obtained through the NIH HIV Reagent Program, Division of AIDS, NIAID, NIH: Anti-Human Immunodeficiency Virus (HIV)-1 gp41/gp120 Monoclonal Antibody (35O22), ARP-12586, contributed by Drs. Jinghe Huang and Mark Connors; monoclonal Anti-Human Immunodeficiency Virus Type 1 (HIV-1) gp120 (2G12), ARP-1476, contributed by Division of AIDS, NIAID; monoclonal Anti-Human Immunodeficiency Virus Type 1 (HIV-1) gp120 Protein, Clone 17b (produced *in vitro*), ARP-4091, contributed by Dr. James E. Robinson. Monoclonal antibody b12 and PGT151 were also produced by Yurogen Biosystems using their in-house human IgG Fc and kappa constant constructs (41, 42). After rigorous washing, cells were lysed in IP lysis buffer (Thermo Scientific, Cat. 87787) and Protease Inhibitor Cocktail (Thermo Scientific, Cat. 78415) according to the manufacturer’s instructions. Virus samples were stored for p24 quantification and concentrated by ultracentrifugation through 6% OptiPrep (Sigma-Aldrich, Cat. D1556). Concentrated virus was lysed under the same conditions as the cell samples.

### p24 enzyme linked immunosorbent assay (ELISA)

The p24 ELISA assay was performed as previously published (43). The capture antibody and standards were obtained from the HIV-1 p24_CA_ Antigen Capture Assay Kit (Frederick National Laboratory) and used as recommended. After incubating samples and standard overnight at 4°C, plates were stained with 1:200 primary rabbit anti-HIV-1 p24 antibody and 1:8000 alkaline phosphatase-conjugated goat anti-rabbit IgG secondary antibody (Jackson Immuno Research Labs, Cat. 111055144). Luminescence was recorded on the Gen5 Microplate Reader and Image Software on the Cytation 3 instrument (BioTek). Nonlinear regression for the standard curve was performed using Prism (GraphPad Software).

### Western blotting

Samples were denatured with Invitrogen NuPAGE LDS Sample Buffer (Fisher Scientific, Cat. NP0007) and 50mM DTT (Thermo Scientific, Cat. A39255) per the manufacturer’s instructions. Following electrophoresis, protein was transferred to a polyvinylidene difluoride (PVDF) membrane using the iBlot 2 Transfer Stacks (Invitrogen, Cat. IB24001) and the iBlot 2 Gel Transfer Device (Invitrogen) and blocked with 5% milk in 0.01% Tween 20 in Tris-buffered saline. gp120 mAb (ARP-1511, Susan Zolla-Pazner, AIDS Reagent Program) was diluted 1:2300 to probe for gp120 Env subunit, mAb Chessie 8 was diluted 1:2000 and p24 Gag antibody (ARP-6457, Michael H. Malim, AIDS Reagent Program) was diluted 1:1000. Peroxidase AffiniPure Donkey Anti-Human IgG (H+L) (Jackson Immuno Research Labs, Cat. 709035149) was diluted 1:50,000 and Goat Anti-Mouse Affinity Purified IgG (H&L) (Rockland, Cat. 61013190500) was diluted 1:6000 to be used as the secondary antibodies. SuperSignal West Femto Maximum Sensitivity Substrate (Thermo Scientific, Cat. 34095) and SynGene Imaging System were used to capture and quantify band signal.

### Ratiometric antibody assay

Jurkat cells were nucleofected with HIV-1 NLCI_QH0692_ using Cell Line Nucleofector Kit V (Lonza, Cat. VCA-1003) and incubated overnight at 37°C. Live cells were enriched with Ficoll-Paque (Cytivia, Cat. 17144003). The cells were stained with 5µg/mL of a fluorescently conjugated antibody (i.e., PGT151, b12 or 2G12) for 45 minutes at 4°C. The antibodies were conjugated to a fluorophore using the Alexa Fluor 488 or 647 Antibody Labeling Kit (Molecular Probes, Cat. A20186 and A20181). After washing, the cells were stained with Live/Dead Fixable Violet (Molecular Probes, Cat. L34964) for 20 minutes at 4°C, followed by a 20-minute fixation with 2% paraformaldehyde (PFA) (Electron Microscopy Sciences, Cat. 15710S) at 4°C. Samples were analyzed using a flow cytometer (Attune NxT Flow Cytometer, Thermo Fisher Scientific). Event-level statistics were performed in R for each antibody (Ab)-pair using data exported from FCS Express Flow Cytometry Software. Events were then grouped for each Ab-pair and within each group, HIV-1-negative and HIV-1-positive cells were downsized to the smaller group size in order to prevent bias due to unequal sample numbers. Outliers were then removed using the interquartile range (IQR) method. To avoid zero or negative values in downstream calculations, the global minimum fluorescence intensity for each Ab was then determined and added to each corresponding group for normalization. Following this, a second round of downsizing was performed to re-equalize group sizes, accounting for imbalances introduced during outlier removal. The following ratio calculations were performed to quantify the shift in Ab binding:

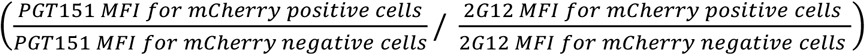

The IQR method was once again applied to remove extreme ratios. The final analysis was graphed, and a statistical analysis was performed.

### Single-round cell-free infection assay and neutralization

Virions were harvested and quantified as described above. Cell-free infection and neutralization assay was adapted from previous publication (12). Briefly, virus containing up to 58ng p24 was incubated with 1×10_5_ RevCEM. After 18 hours, the media was replaced with complete RPMI media with 10μM azidothymidine (AZT) (NIH HIV Reagent Program, NIAID, HRP-3485) to ensure infection was limited to a single round. After 40 hours, cells were washed and stained with Live/Dead Fixable Violet or Green Dead Cell Stain Kit (Molecular Probes, Cat. L34964 and L23101). Cells were then fixed in 2% PFA and analyzed by flow cytometry. Negative live/dead stain and positive mCherry signal from the cell population represent alive and infected cells. In neutralization studies, the virus was pre-incubated with monoclonal antibodies (mAbs) for 30 minutes at 37°C prior to culturing with cells. As a negative control, anti-CD4 antibody Leu3a (BC Cell Analysis, Cat. 3P340853) was incubated with cells.

### Cell-to-cell infection assay and neutralization

Cell-to-cell infection neutralization assay was adapted from a previous publication (40). Env expression in Jurkat T cells (donor) was generated as described above. Donor and target cells were dye-labeled with cell proliferation dye eFluor 670 and eFluor 450, respectively (Invitrogen, Cat. 65084090 and 65084290). 1×10^5^ donor cells were co-cultured with equal amounts of target cells. After 18 hours, the media was replaced with RPMI complete and 10μM AZT. After 40 hours, the cells were washed and stained with a live/dead cell stain. After washing, cells were fixed with 2% PFA for 20 minutes at room temperature and analyzed by flow cytometry. Positive mCherry signal from eFluor 450 cell population represents target cells that became HIV-1 infected. The neutralization assay was conducted by incubating donor or target cells with mAbs for 30 minutes at 37°C prior to co-culture.

### Cell-to-cell transfer assay and transfer-neutralization

Cell-to-cell transfer and transfer-neutralization assay were performed as previously described (40). The preparation of donor and target cells is as described for the cell-to-cell infection assay with HIV-1 Gag-iCherry clones. After co-culturing for 6 hours, cells were washed and trypsinized to remove surface-attached free virions and neutralized with complete RPMI media. The positive mCherry signal from the eFluor 450 cell population represents target cells that received transferred viral particles. To perform neutralization, the mAbs were incubated with the donor or target cells for 30 minutes at 30°C before co-culturing. The samples were analyzed by flow cytometry.

## Results

### Comparison of endocytic-and cleavage-deficient Env mutants

Mutating Env’s main endocytic motif can alter the neutralization potency and efficacy of cell-to-cell infection in an epitope-dependent manner, suggesting that endocytosis may influence Env’s antigenic state (12). Since Env cleavage can affect antibody recognition by altering epitope exposure and glycan composition (18–22), and because endocytic recycling pathways are implicated in the packaging of Env onto virions (44–48), we examined whether eliminating Env’s dominant endocytic motif impacts its cleavage status on cells and virions. To generate an endocytic-deficient Env (ASPI-Env) of the T/F HIV-1 QH0692 clone, we substituted the tyrosine in the membrane-proximal motif at 724 with alanine (Y724A) (**Fig. 1A**). As our hypothesis was focused on observing cleavage-dependent differences, we also generated an uncleavable Env mutant (SEKS-Env) by mutating the furin cleavage site at the gp120-gp41 junction, _519_REKR_522_, to SEKS (**Fig. 1A**). These mutations were introduced into the cherry-fluorescent protein expressing HIV-1 clone, NLCI (**Fig. 1B**) (38), and a viral clone that produces fluorescent Gag-iCherry HIV-1 particles (**Fig. 1C**) (39).

**Figure 1.**
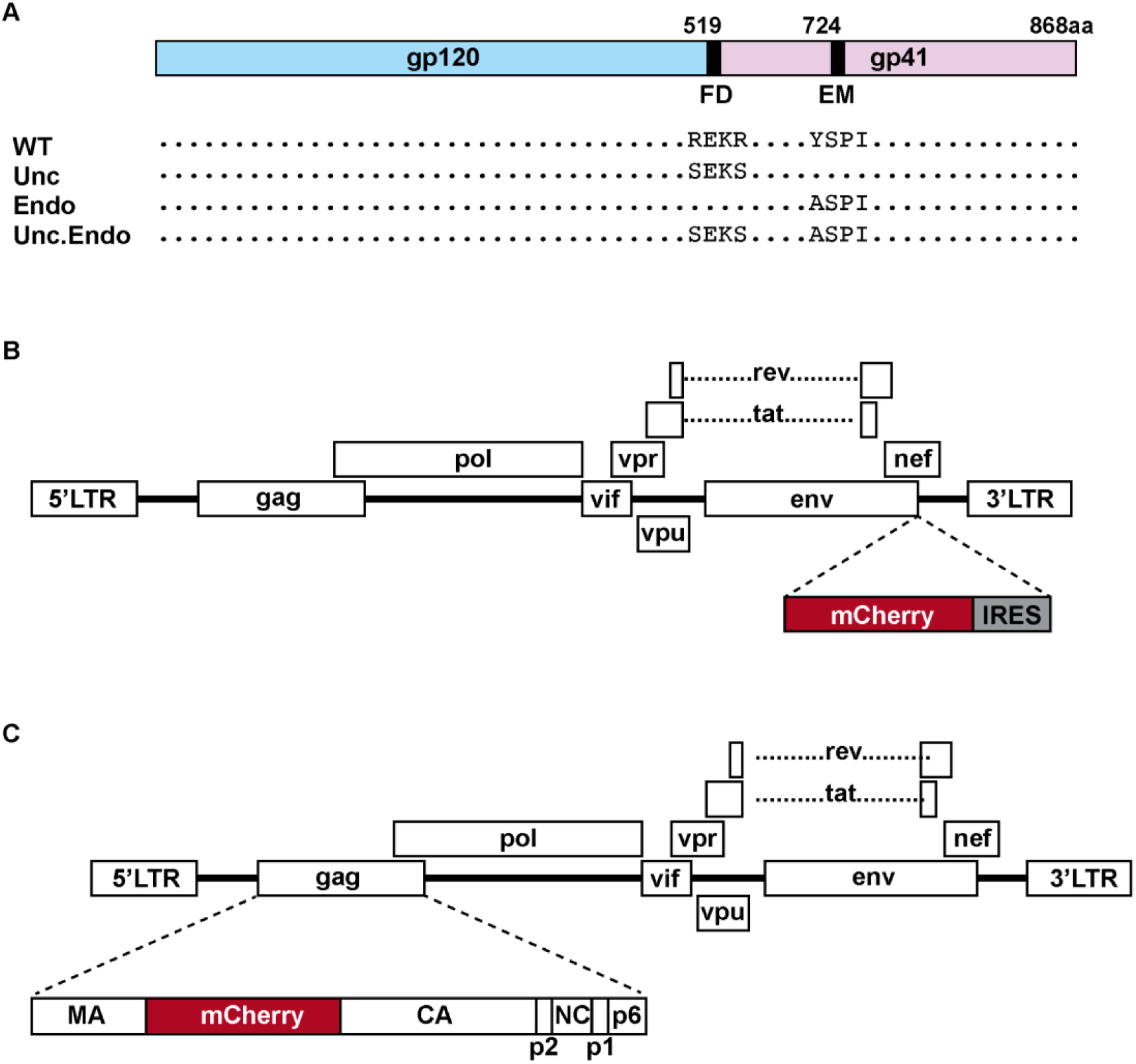
Endocytosis and furin cleavage site mutations within fluorescent reporter HIV-1. **(A)** Schematic representation of Env mutations in the furin cleavage site (FS) and main endocytic motif (EM). Proviral DNA of T/F QH0692 ENV was cloned into the NL43 backbone, producing an uncleavable (Unc), endocytosis-deficient (Endo), and double mutant (Unc.Endo) variants. **(B)** NLCI construct includes an mCherry and internal ribosome entry site (IRES) upstream of the nef gene. **(C)** GagiCherry construct consists of an mCherry signal between the matrix (MA) and capsid (CA) genes.

### Env glycosylation and antibody recognition on cells and virions of endocytic-and cleavage-deficient Env mutants

In prior studies, we and others have shown that disruption of the main endocytic motif decreased infectivity (30, 48, 49) and modestly enhanced the incorporation of uncleaved Env into 293T-derived virions (12, 49). To investigate how the cleavage status of Env, present in cell or virus lysates, is altered by mutating the endocytic motif, or the furin cleavage motif, or both, in T/F QH0692 Env, we expressed wildtype and mutant NLCI_QH0692_ in 293T cells. The cells and viruses were collected for western blot analysis to examine the relative abundance and glycan composition of bands that represent cleaved and uncleaved Env. Antibodies and lectins were used to probe the whole-cell and viral lysates **(Fig. 2A)**. In whole-cell lysates, three bands can be detected in WT- and ASPI-Env expressing cells, while SEKS- and SEKS.ASPI-Env expressing cells show only two (**Fig. 2B**). A gp41 antibody, Chessie 8, detects the two highest molecular weight bands, consistent with them representing two isoforms of uncleaved Env (gp160) (**Fig. 2B**). The bottom band, which is not reactive with Chessie, is cleaved Env (gp120) (**Fig. 2B**).

**Figure 2.**
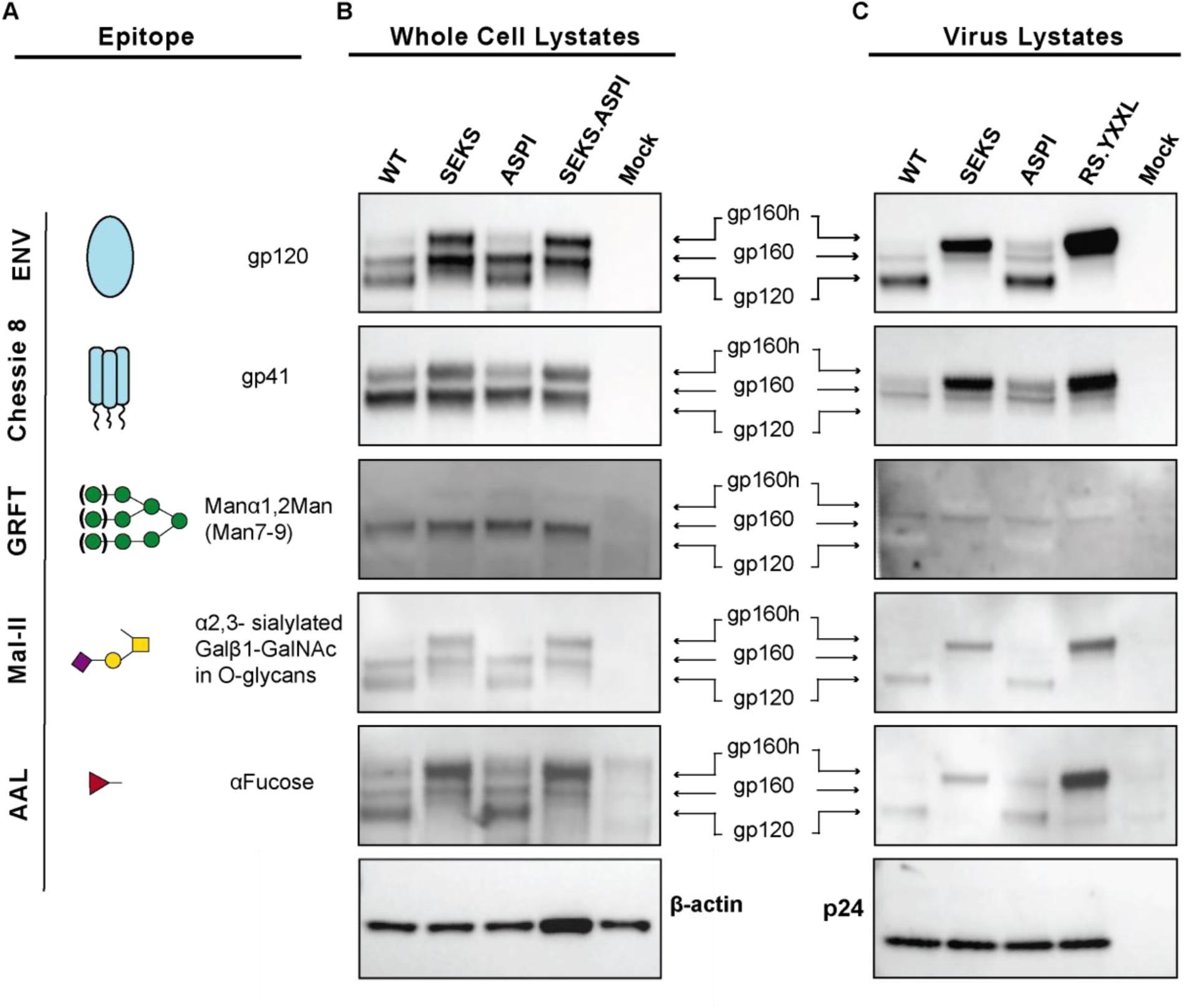
Expression of cleaved and uncleaved glycoforms of Env within the cell and on virions, and the impact of ASPI mutation in comparison to the uncleavable SEKS mutant. **(A)** Epitope of anti-Env Abs and lectins used for Western blotting. WT and mutant NLCI_QH0692_ were expressed in **(A)** 293T cells and **(B)** concentrated viral particles. Detection was performed using antibodies against gp120 (268-D IV), gp41 (Chessie 8) and lectins (GRFT, Mal-II, AAL). β-actin and p24 were probed as loading controls for whole-cell lysates and virus lysates, respectively.

To determine whether the two gp160 bands are glycovariants, we blotted with a panel of three lectins. Griffithsin (GRFT), which binds to high-mannose (50, 51), only bound to the middle Env band in whole-cell lysates and viral lysates (**Fig. 2B-C**). Notably, GRFT detection in the viral lysates is poor, as none of the WT or mutant Envs had a high concentration of the middle, high-mannose gp160 glycoform. In comparison, *Maackia amurensis* lectin II (MAL II) and *Aleuria aurantia* lectin (AAL), which bind to sialylated (52) and fucose glycans (53), respectively, both bound to all three Env bands (**Fig. 2B-C**). The difference in lectin binding observed within the gp160 bands suggests that the higher-molecular weight band is a differentially glycosylated form of uncleaved Env, which we refer to as gp160h, and the lower band as gp160.

Env glycosylation is influenced by its cleavage status, with cleaved Env enriched in oligomannose glycans, while uncleaved Env carries a higher proportion of complex glycans (22). This may explain the two gp160 glycovariants observed in the SEKS- and SEKS.ASPI-Env, as the increased conformational flexibility of uncleaved Env likely permits alternative glycan processing. Additionally, AAL showed greater binding to gp120 and gp160h (**Fig 2B**). We observed a similar glycosylation profile of gp160h and gp120, which were both selectively packaged onto the viral particle with greater abundance than gp160. A small increase in gp160h can also be observed in the viral lysates for ASPI-Env-derived particles compared to WT-Env particles (**Fig. 2C**). Due to the low abundance of gp160 on viral particles, replicates could not confirm a significant increase in gp160h.

### Endocytic motif reduces cell surface expression of cleaved Env

Next, we investigated whether the motif altered the engagement of antibodies whose epitope exposure is influenced by Env’s cleavage status. We performed a cell-surface immunoprecipitation experiment by adapting the pulldown assay described by Zou et al. (28) in which antibodies are bound to cell surface Env, and excess antibody is washed off prior to cell lysis and pulldown. mAbs were selected for the pulldown based on existing research regarding the mAbs’ preference for binding to either cleaved or uncleaved Env, highlighted in **Table 1**. PGT151 is a mAb that targets a quaternary structure spanning the gp120-gp41 interface and is highly specific for cleaved Env (42, 54). In contrast, 35O22 also targets the gp120-gp41 interface, but at a distinct site, and, unlike PGT151, is not cleavage-dependent, binding both cleaved and uncleaved Env (55). Similarly, b12 is a CD4-binding site antibody that has been shown to bind both cleaved and uncleaved Env; however, several studies indicate it has a moderate preference for uncleaved Env (19, 20, 54, 56). Finally, 2G12 targets an oligomannose-based epitope and is therefore regarded as conformationally independent in the literature (19, 20, 57, 58).

**Table 1.**
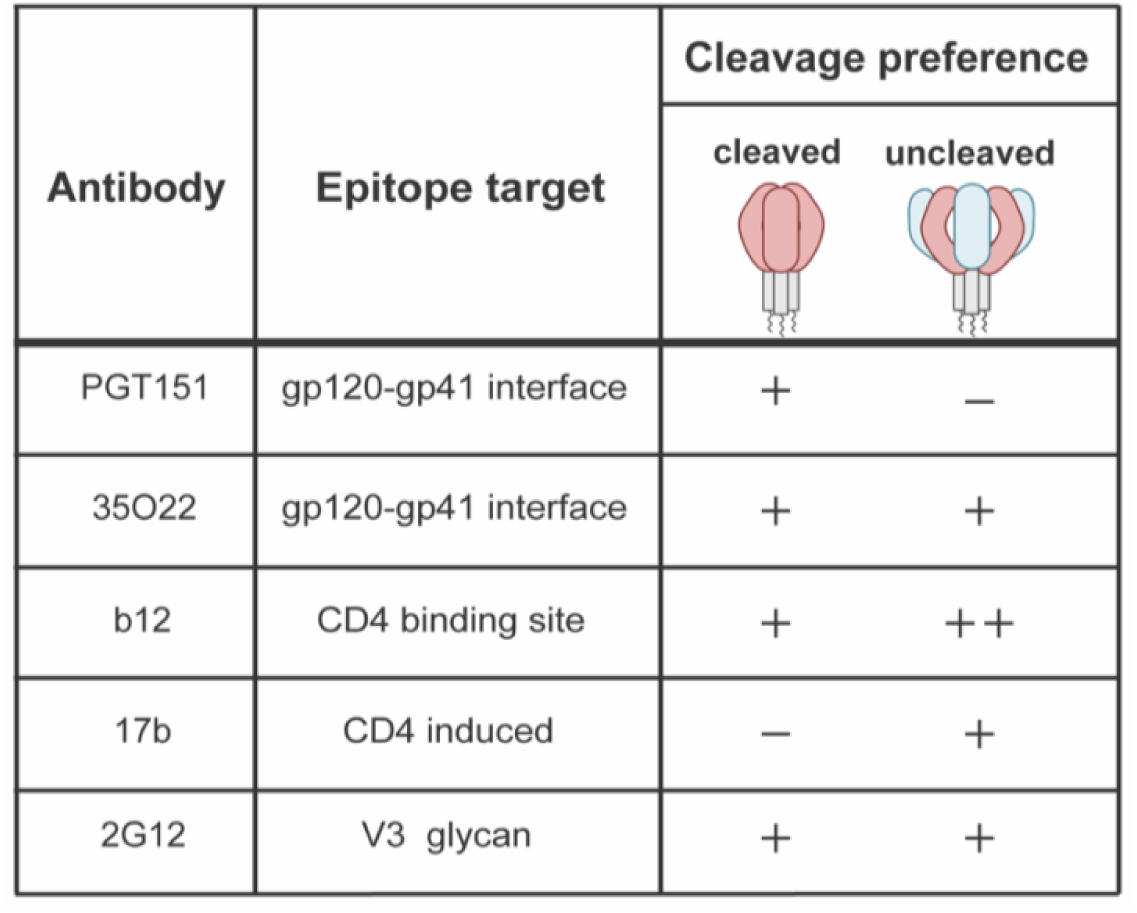
Epitopes of bNAbs that target cleavage-sensitive epitopes.

| Antibody | Epitope target | Cleavage preference |  |
| --- | --- | --- | --- |
|  |  | cleaved | uncleaved |
| PGT151 | gp120-gp41 interface | + | — |
| 35O22 | gp120-gp41 interface | + | + |
| b12 | CD4 binding site | + | ++ |
| 17b | CD4 induced | — | + |
| 2G12 | V3 glycan | + | + |

The IP of cell-surface WT-Env indicates that there is a heterogeneous mixture of cleaved and uncleaved Env present on the cell surface (**Fig. 3A**). The 35O22 pulldown highlights this as both gp160 and gp120 can be detected in cells expressing WT- and ASPI-Env. In comparison, PGT151 preferentially pulled down gp120, while b12 predominantly pulled down gp160. 2G12 also showed a preference for targeting gp160 (**Fig. 3A**). A decrease in gp120 pulldown from PGT151 is observed in the ASPI-Env expressing cells compared to WT-Env, consistent with a reduction in cleaved Env at the cell surface in this mutant (**Fig. 3A**). To quantify the loss of cleaved Env with ASPI-Env, we performed densitometry analysis of the gp120 band from PGT151 relative to the gp160 band from 2G12 pulldown (**Fig. 3B**). Quantification confirmed a decrease in the cleavage ratio of ASPI-Env compared to WT-Env from the cell surface (**Fig. 3B**). This decrease was consistent with the loss of cleavage observed in the cell lysates quantified between ASPI-Env and WT-Env expressing cells (**Fig. 3C**). These data support that disrupting Env endocytosis reduces cleaved Env on the surface of infected cells.

**Figure 3.**
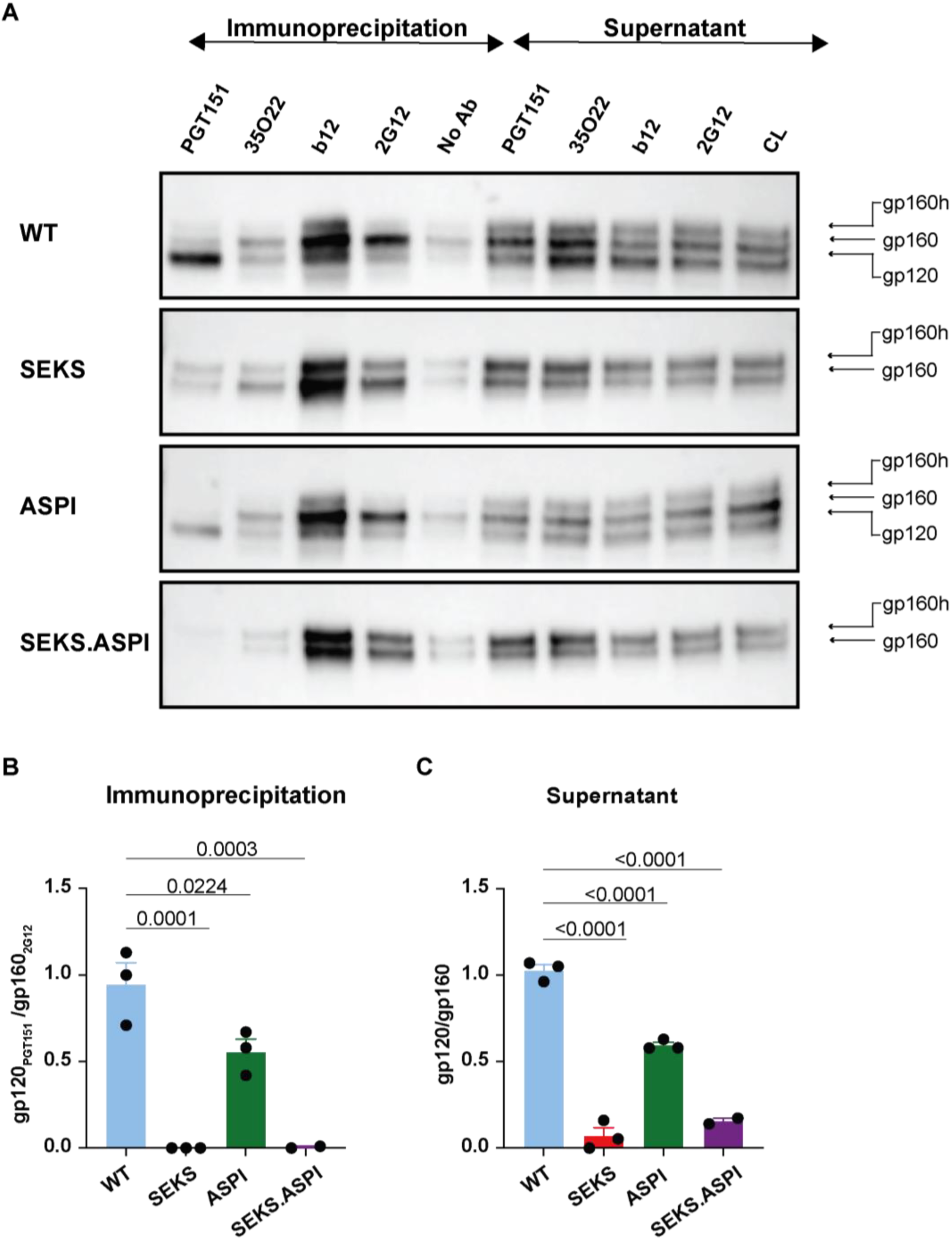
Immunoprecipitation of cell-surface Env with Abs that target cleavage-sensitive epitopes selectively recognizes distinct isoforms of Env. **(A)** Cell surface Env was immunoprecipitated from 293T cells transfected with WT and mutant NLCI_QH0692._ Immunoprecipitation was performed with mAbs (PGT151, 35O22, b12, and 2G12). Western blotting was performed with anti-Env antibody gp120 (268-D IV). **(B)** Densitometry analysis for gp120 band (by PGT151) and gp160 band (by 2G12) provides a cleavage ratio of cell-surface Env. **(C)** Total Env cleavage ratio is calculated by performing densitometry analysis on gp120 and gp160 band in the cell lysates (CL). Densitometry data from three independent experiments with SEM are shown. Dunnett’s multiple comparisons test p-values shown.

As expected, the SEKS mutation, which fully abrogates cleavage, was not detected in the cell surface IP by PGT151 in either SEKS- or SEKS.ASPI-Env (**Fig. 3A**). IP with 35O22, b12, and 2G12 all captured both gp160 and gp160h in SEKS- and SEKS.ASPI-Env (**Fig. 3A**). This reveals that the selective sorting of gp160h over gp160 onto virions in **Figure 2C** is not a result of ineffective trafficking of gp160 to the cell surface. Rather, the pulldown indicates that while both glycoforms of Env are present on the cell surface, only one is selected for packaging. This data suggests that glycosylation may influence Env packaging.

### Ratiometric antibody binding reveals increased uncleaved:cleaved ratio in Env endocytic mutant on the cell surface

To assess how Env endocytosis can affect the conformation and cleavage status of Env present on the T cell surface, we expressed WT and mutant HIV NLCI_QH0692_ in Jurkat T cells and measured binding to the panel of mAbs discussed above. Previous literature has found that disrupting Env’s main endocytic motif enhances cell surface Env expression (12, 59–61). Therefore, staining with only one mAb would not inform whether changes to Ab binding were a result of increased Env on the cell surface due to the ASPI mutation, or changes to Env’s conformation that increase or decrease exposure to the Abs’ epitope. To address this, we performed a ratiometric antibody binding assay that uses two-color flow cytometry to measure the median fluorescence intensity (MFI) of an Ab that targets cleavage-sensitive epitopes (i.e., PGT151 or b12) relative to the MFI of an Ab whose epitope exposure is considered independent of cleavage status (i.e., 2G12). Antibodies were added simultaneously, as sequential binding, particularly with 2G12 added first, disrupted binding and was inconsistent with single Ab staining (data not shown).

We employed a ratiometric fluorescence assay to compare mCherry_+_ (HIV-1_+_) cells expressing WT and mutant Env **(Fig. 4)**. Compared to WT-Env, ASPI-Env showed a loss of PGT151 binding relative to 2G12 binding (**Fig. 4A**). A similar shift, but with greater magnitude is observed when comparing WT-Env to SEKS- and SEKS.ASPI-Env (**Fig. 4A**). The relative decrease in PGT151 binding is consistent with the requirement for cleavage to stabilize the pre-fusion conformation necessary for binding (62). This indicates that disrupting Env internalization reduces the exposure of the cleavage-dependent epitope on cell-surface Env.

**Figure 4.**
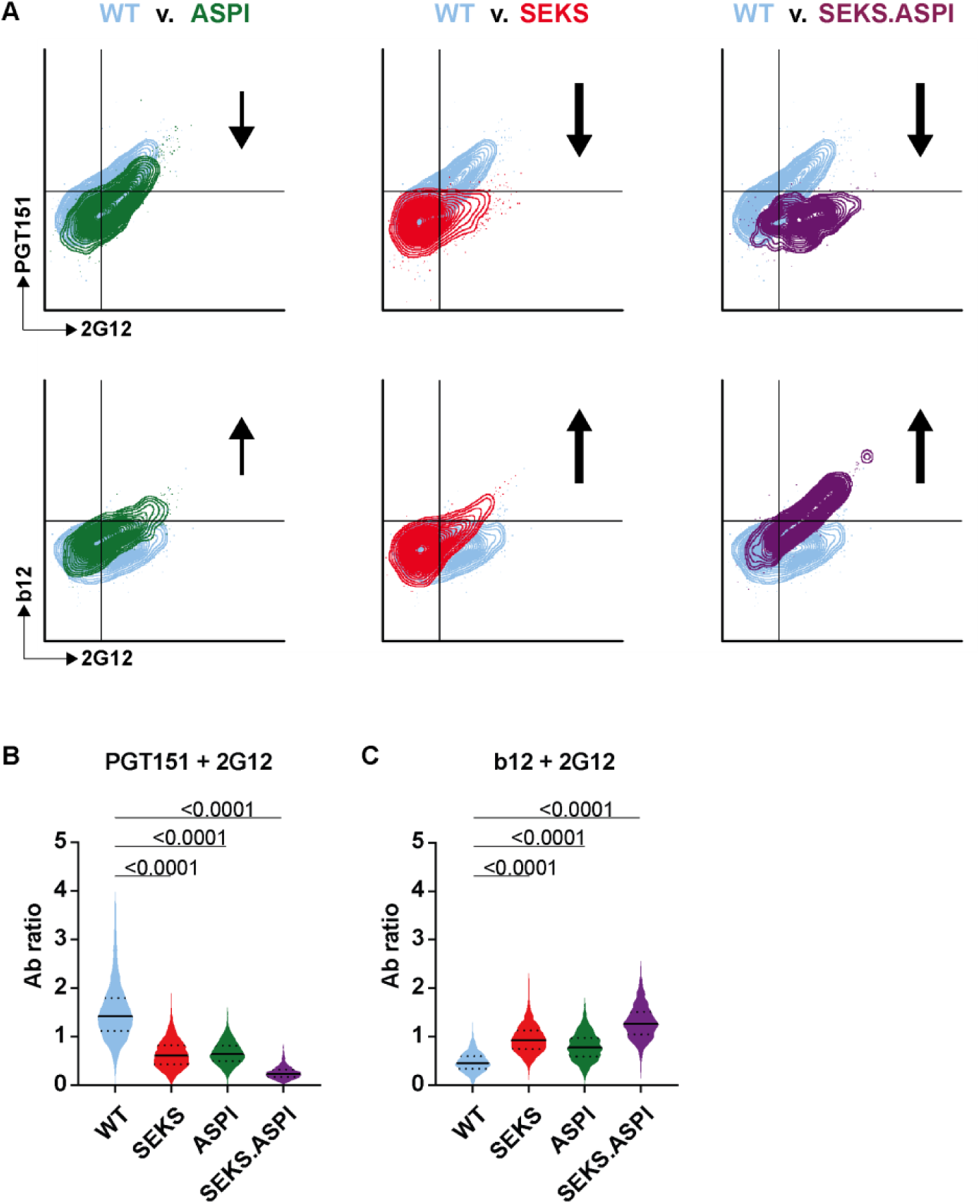
Dual fluorescence flow cytometry assay of Env with Abs that target cleavage-sensitive epitopes. **(A)** Representative flow cytometry dot plots show Ab staining of cell surface WT and mutant NLCI_QH0692_ in Jurkat T cells. Arrows represent the Ab binding shift. **(B-C)** The antibody binding ratio was calculated as the fluorescence intensity for each Ab pair in the HIV-positive over the HIV-negative population of cells. Event-level statistics were conducted across four independent experiments, as shown. Dunnett’s multiple comparisons test p-values are shown.

To study changes to Env’s conformation indicative of a more open conformation, we also stained with b12, a mAb which in several studies has displayed a moderate affinity for uncleaved Env relative to cleaved Env (19, 20, 54, 56), relative to 2G12. As **Fig. 4A** demonstrates, ASPI-Env exhibited enhanced binding to b12 over 2G12 compared to WT-Env, also mirroring shifts with SEKS- and SEKS.ASPI-Env. Note that although the proportion of WT-Env expressing cells binding b12 appears low in the ratiometric assay, it is consistent with the percentage of b12-positive cells observed in single antibody staining (data not shown). All mutants demonstrated significant loss of PGT151 binding relative to WT-Env expressed on the cell surface (**Fig. 4B**). In comparison, the shift in b12 binding relative to 2G12 was significantly enhanced between WT and mutant Env (**Fig. 4C**). These data suggest that mutating the endocytic motif has altered the antigenic profile of cell surface Env and increased binding was not simply due to enhanced cell surface expression. Moreover, the shifts in binding pattern observed with the ASPI mutant mimic shifts observed with the SEKS mutant. We conclude that the ASPI mutation enhances the presentation of epitopes that are preferentially exposed in uncleaved Env.

### Endocytic motif mutation also enhances the exposure of epitopes associated with uncleaved Env on virions

To investigate the impact of an endocytic-deficient Env on HIV-1 infectivity and sensitivity to neutralizing antibodies, we generated cell-free wild-type and mutant viruses (NLCI_WT_, NLCI_SEKS_, and NLCI_ASPI_) from transfected 293T cells. mCherry is expressed in place of nef, allowing for the detection of active HIV-1 gene expression (**Fig. 1B**). The viruses were titrated onto CCR5-expressing RevCEM cells. Viral particles with ASPI-Env were infectious, consistent with previous findings (**Fig. 5A**) (30, 48, 49). No infection was also observed with NLCI_SEKS_ (**Fig. 5A**). This also aligns with prior studies, which show that without cleavage, Env is not fusogenic and therefore non-infectious (62).

**Figure 5.**
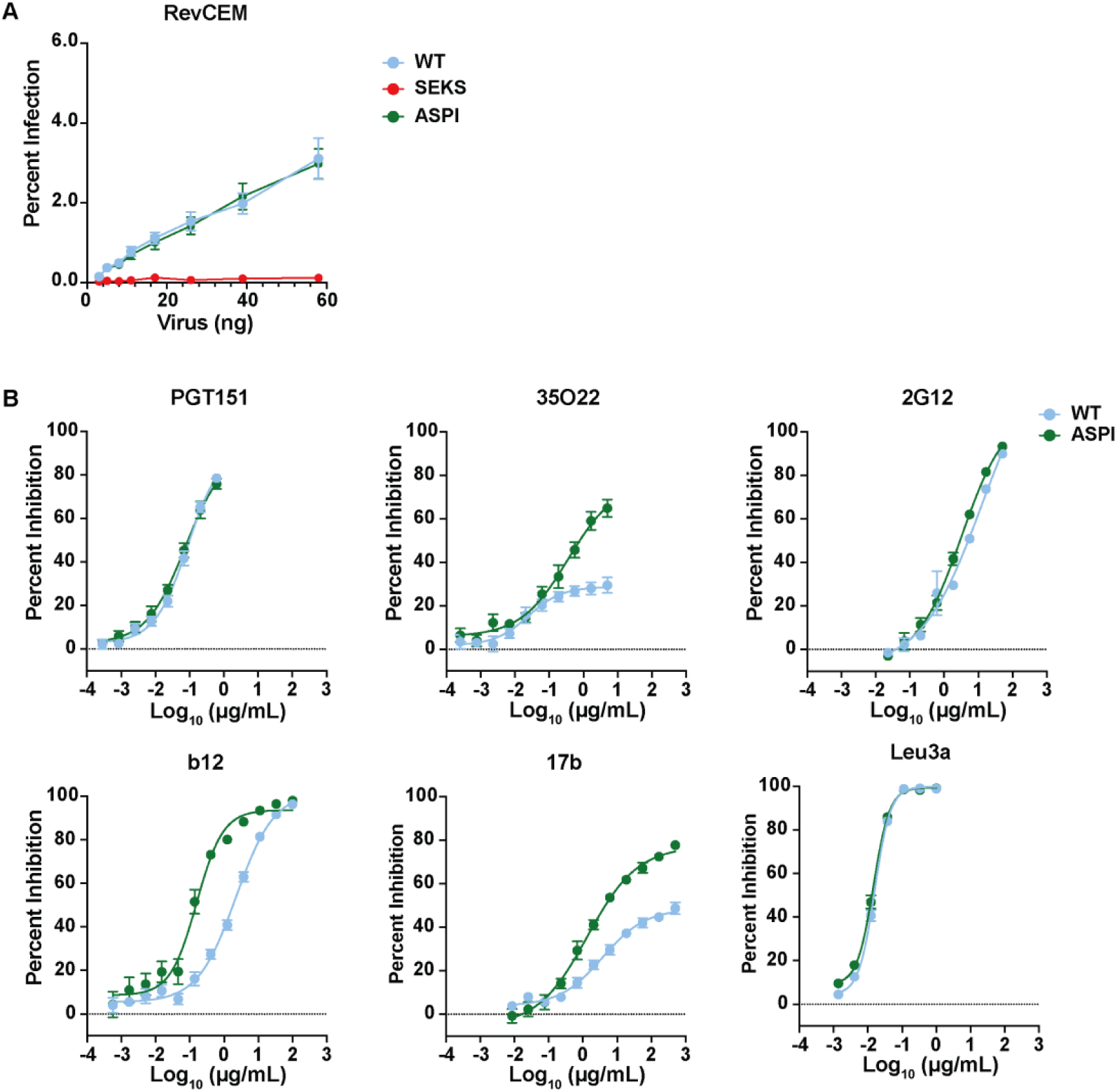
Neutralization assays of cell-free infection comparing WT vs endocytic motif mutant using Abs that target cleavage-sensitive epitopes. **(A)** WT and mutant NLCI_QH0692_ cell-free virions produced in 293T cells were titrated onto RevCEM cells. p24 ELISA was used to determine virus concentration. **(B)** Neutralization of cell-free infection against WT and ASPI NLCI_QH0692_ by PGT151, 35O22, 2G12, b12, 17b, and Leu3a. The virus was incubated with mAbs for 30 minutes at 37°C before culturing with cells. The mean and SEM data are shown for at least three independent experiments.

Next, we performed neutralization with the panel of mAbs previously discussed. Additionally, to better inform on how Env’s conformation changes upon cleavage status, we also tested 17b, mAb that binds to an epitope exposed upon Env-CD4 binding and which is found in open Env conformations independent of CD4 binding (63–65). (**Table 1**). As a control, neutralization was also performed using the CD4-binding antibody Leu3a. Leu3a inhibits HIV-1 infection by competing with Env for CD4 binding and entry. No difference was observed in Leu3a’s sensitivity to NLCI_WT_ and NLCI_ASPI_ (**Fig. 5C**). No difference was also observed in the sensitivity of NLCI_WT_ and NLCI_ASPI_ to neutralization by PGT151 and 2G12 (**Fig. 5C**). In contrast, NLCI_ASPI_ showed enhanced sensitivity to neutralization by 35O22, b12, and 17b (**Fig. 5C**), indicating greater access to epitopes exposed on uncleaved Envs. At the highest concentration tested for 35O22 (10μg/mL), only ∼20% of NLCI_WT_ was neutralized (**Fig. 5C**). By comparison, at the same concentration, 35O22 neutralized ∼60% of NLCI_ASPI_ (**Fig. 5C**). Similarly, 17b neutralized ∼40% of NLCI_WT_ compared to ∼80% of NLCI_ASPI_ at the highest concentration tested (1000μg/mL) (**Fig. 5C**). In the case of b12, 100% of NLCI_WT_ and NLCI_ASPI_ was neutralized (**Fig. 5C**). However, b12 showed enhanced potency against NLCI_ASPI,_ with its half-maximal inhibitor concentration (IC_50_) being 10-fold less than that of NLCI_WT_ (**Fig. 5C**). The increased sensitivity of NLCI_ASPI_ to 35O22, 17b, and b12 is striking, suggesting that small increases in gp160 Env packaging can alter neutralization of cell-free HIV-1 infection.

### Endocytic motif mutation enhances b12 sensitivity to cell-to-cell infection

Having observed that mutating Env’s endocytic motif increased uncleaved Env on the cell surface, we next investigated how cell-to-cell infection and neutralization would be impacted. The cell-to-cell infection assay was performed as in Durham et al. (40). Briefly, Jurkat T cells (donor cells) nucleofected with wild-type and mutant NLCI were co-cultured with RevCEM cells (target cells). The donor and target cells were dye-labeled to detect infection in the target cells, allowing distinction through flow cytometry. To ensure that differences in infectivity could be attributed to the endocytic and cleavage site mutations and not to differences in nucleofection efficiency, the amount of mCherry expression in the cells was measured prior to co-culturing the cells. To ensure equal nucleofection efficiency, untreated donor cells were spiked into the wildtype- and mutant-expressing donor cells. NLCI_SEKS_ is not infectious and was therefore not tested. Both NLCI_WT_ and NLCI_ASPI_ were infectious through the cell-to-cell assay (**Fig. 6A**). Next, we tested the sensitivity of the mAbs discussed in the previous section to inhibit NLCI_WT_ and NLCI_ASPI_ cells. When compared to NLCI_WT_, NLCI_ASPI_ cell-to-cell infection did not exhibit increased sensitivity or resistance to PGT151, 35O22, 2G12 or 17b (**Fig. 6C**). Notably, 35O22 showed no activity against either NLCI_WT_ or NLCI_ASPI_ cell-to-cell infection, despite demonstrating inhibition against cell-free virus (**Fig. 6C** **and** **Fig. 5C**). NLCI_ASPI_ cell-to-cell infection was, however, dramatically more sensitive to b12 neutralization and shifted the non-sigmoidal inhibition curve of NLCI_WT_ into a more sigmoidal pattern (**Fig. 6C**). To neutralize 80% of NLCI_WT_, b12 required ∼143μg/mL (**Fig. 6C**). In contrast, neutralization of NLCI_ASPI_ was achieved with approximately 0.5μg/mL (**Fig. 6C**). b12’s unusually broad neutralization curve against WT-Env appears biphasic, suggesting there are two populations of Env engaging in cell-to-cell infection. We also observed a difference in the neutralization of NLCI_WT_ and NLCI_ASPI_ cell-to-cell infection by Leu3a at concentrations above 0.1μg/mL (**Fig. 6C**). The neutralization curves demonstrate that although the amount of uncleaved Env expressed on the cell surface due to the ASPI mutation increased, only b12 showed enhanced sensitivity to inhibition of cell-to-cell infection. This suggests that disrupting endocytosis has large effects only in a specific subset of antibodies. The changes in b12’s neutralization curve from a multiphasic curve into a sigmoidal curve suggest that multiple conformations of Env are facilitating cell-to-cell infection in the WT Env, whereas the sigmoidal curve against ASPI-Env increases the uniformity of Env targeted.

**Figure 6.**
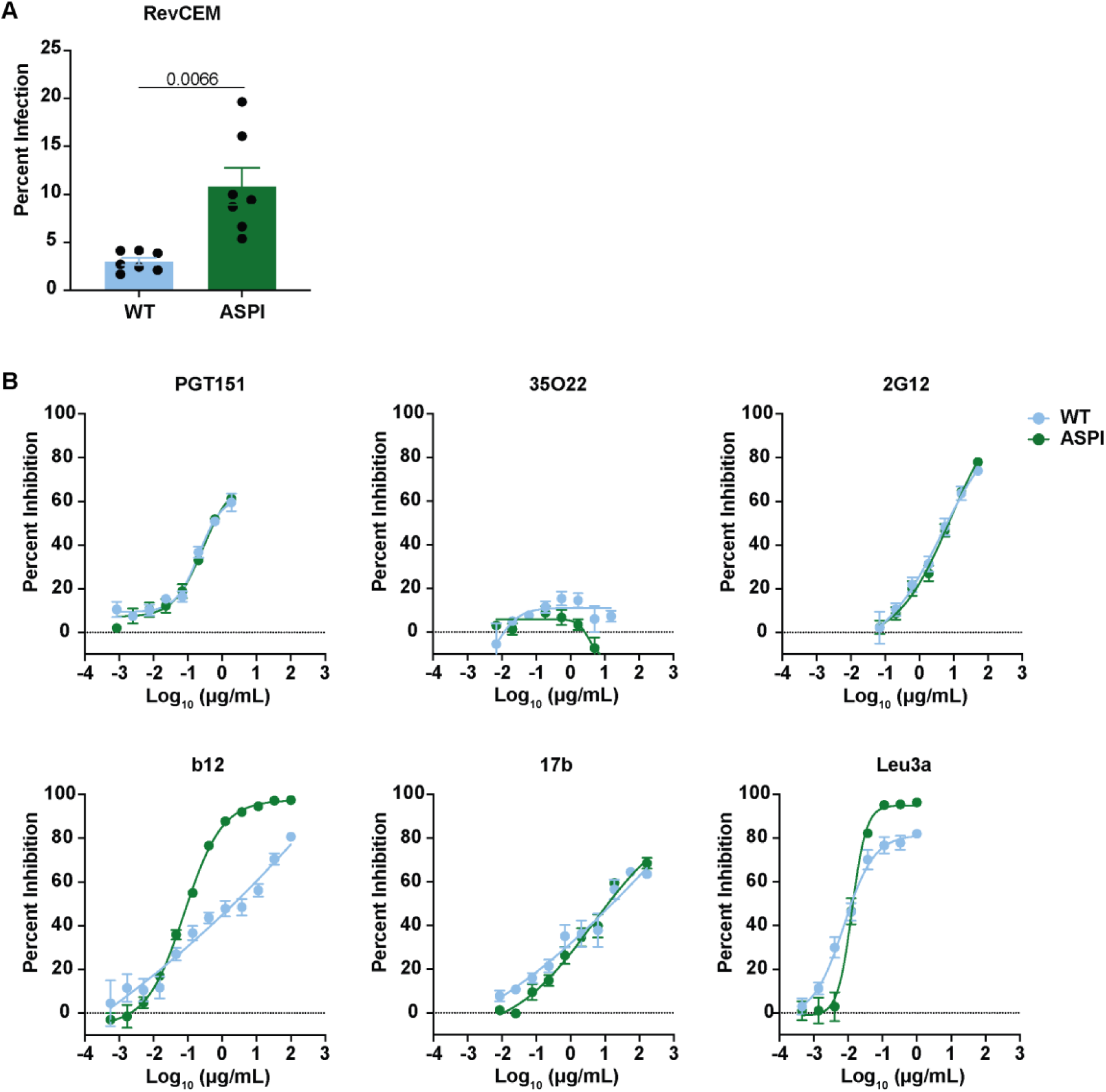
Neutralization assays of cell-to-cell infection with Abs that differentially recognize cleaved versus uncleaved Env. **(A)** Nucleofected Jurkat T cells expressing WT and ASPI NLCI_QH0692_ were cocultured with RevCEM cells. Nucleofection efficiency was determined by measuring mCherry expression in Jurkat T cells 24hrs after nucleofection using flow cytometry. Nucleofection efficiency was normalized by spiking in mock nucleofected Jurkat T cells prior to co-culturing. p-value for Unpaired t test with Welch’s correction and unpaired t test are shown, respectively. **(B)** Neutralization of WT and ASPI NLCI_QH0692_ cell-to-cell infection by PGT151, 35O22, 2G12, b12, 17b, and Leu3a. The donor cells were incubated with anti-Env mAbs and target cells with Leu3a for 30 minutes at 37°C before co-culturing cells. All experiments shown represent at least three independent experiments. The mean and SEM data are shown for at least three independent experiments.

### Env endocytic motif influences lectin sensitivity to HIV-1 infection

Dense glycosylation of Env is an important property of HIV-1’s defense against antibodies (66–68). Because Env glycosylation can vary according to cleavage state (22), we may expect that changes in Env’s cleavage state will also modulate glycan composition, as cleavage limits access to Golgi glycosidases. Many bNAbs have also been found to be glycan-dependent (23–27). This warrants further exploration of how the main endocytic motif of Env affects its glycan composition, particularly Env engaged in HIV-1 infection. To address this, we performed cell-free and cell-to-cell neutralization assays as described above, using lectins instead of monoclonal antibodies (mAbs). The lectins, highlighted in **Figure 2A**, have been described in the literature to bind glycans with high-mannose (GRFT) (50, 51), sialylation (Mal-II and SLBRN) (52, 69), and fucose (AAL) (53). GRFT, in particular, has been observed to have antiviral activity against a range of pathogens, including simian immunodeficiency virus (SIV) and hepatitis C virus (HCV) (70–72). GRFT demonstrated potent inhibition of cell-free and cell-to-cell WT infection, with an IC_50_ of 0.01-0.002μg/mL (**Fig. 7A, B**). In comparison, Mal-II, AAL, and SLBRN all demonstrated either enhanced or no effect on WT cell-free and cell-to-cell infection.

**Figure 7.**
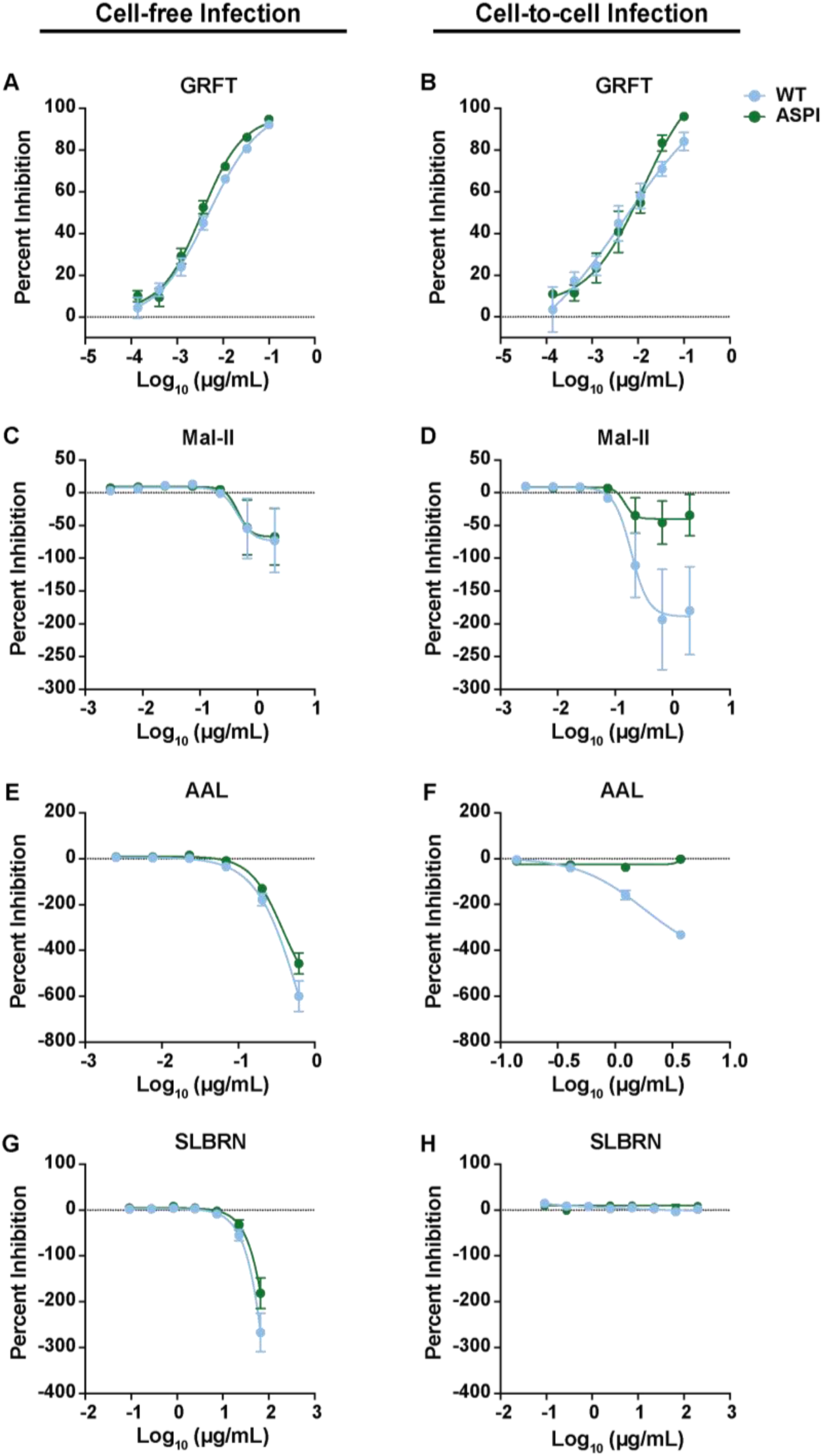
ASPI mutation reduces the enhancement activity of two lectins in cell-to-cell infection. Cell-free and cell-to-cell neutralization assays were performed as discussed above with RevCEM as target cells. **(A-B)** GRFT, **(C-D)** Mal-II, **(E-F)** AAL, and **(G-H)** SLBRN lectins were incubated with cell-free viral particles or nucleofected Jurkat T cells at 37 °C for 30 minutes before adding target cells. In both assays, cultures were treated with AZT 18 hours post-infection. The mean and SEM data are shown for at least two independent experiments.

Our study revealed that the potency of each lectin depended on the mode of HIV-1 infection. Specifically, Mall-II enhanced cell-to-cell infection by ∼3-fold, while cell-free infection was enhanced by ∼1.5-fold at a concentration of ∼1μg/mL (**Fig. 7C, D**). In comparison, AAL was more effective at enhancing cell-free infection, showing a 7-fold increase compared to a 4-fold enhancement in cell-to-cell infection at 3.7μg/mL (**Fig. 7E, F**). The loss of cell-to-cell infection activity was also observed with SLBRN, a lectin that binds sialylated O-glycans (**Fig. 7G, H**) (69). These differences in lectin activity in cell-free and cell-to-cell infection suggest that the glycan composition of Envs engaged in each mechanism of infection differs.

The data also demonstrate that disrupting the endocytic motif affected the lectin’s activity. While cell-free infection remained immune to changes in Env’s endocytic motif (**Fig. 7A, C, E, G**), Mall-II and AAL exhibited reduced activity against the ASPI mutant compared to WT in cell-to-cell infection (**Fig. 7D, F)**. However, this effect was lectin-dependent as ASPI-Env did not affect the inhibitory effect of GRFT or the inactivity of SLBRN in cell-to-cell infection (**Fig. 7B, H**). These data further suggest that Env engaged in cell-free and cell-to-cell infection is antigenically distinct, with glycosylation likely contributing to these differences, and that endocytosis of Env on the cell surface may contribute to the conformational state and glycosylation composition of Env engaged in these two modes of infection.

### Investigating the function of uncleaved Env during cell-to-cell infection

Uncleaved Env is not fusogenic and, consequently, is not thought to participate in HIV-1 infection (62, 73). However, studies have shown that uncleaved Env can bind to CD4 (16, 74). Considering the increase in uncleaved Env observed with the ASPI mutation, we investigated whether uncleaved Env on the cell surface can participate in the first CD4-dependent step in cell-to-cell infection, the transfer of virions from infected donor to uninfected target cells. Specifically, we were interested in determining whether uncleaved Env could mediate the transfer of HIV-1 particles between cells by initiating the formation of a VS by binding to CD4. To test this, we performed a cell-to-cell transfer assay similar to the cell-to-cell infection assay described above, with a few notable differences. First, we used an HIV-1 construct that expresses mCherry-tagged Gag particles (**Fig. 1C**). This enables us to detect the transfer of fluorescent viral particles as they are taken up by target cells. Before collection, the cells are treated with trypsin to remove any surface-bound viral particles that could produce a false-positive transfer. Furthermore, the viral transfer assay is limited to six hours, as replication is attenuated, ensuring we are measuring only CD4-dependent viral transfer and not infection. To confirm the transfer of the fluorescent viral particles is CD4-dependent, target cells are incubated with 1μg/mL of Leu3a prior to co-culturing. Importantly, Leu3a binds to the CD4 receptor, thereby inhibiting CD4-Env binding and consequently both VS formation and viral fusion. When coculturing T cells expressing WT-Env or ASPI-Env, both showed reproducible transfer of virions to RevCEM cells **(Fig. 8A)**. Importantly, transfer of the virus particle decreases upon treatment with Leu3a, thereby demonstrating the transfer observed is CD4-dependent **(Fig. 8A)**. The cell-to-cell transfer of infected cells expressing SEKS-Env also showed a significant increase in CD4-dependent transfer of viral particles compared to WT-Env (**Fig. 8A**). As SEKS-Env is not fusogenic, viral and target cell membrane fusion is not expected. However, viral particles at cell-contact sites have been observed to undergo endocytosis (4, 8, 75, 76). This indicates that although SEKS-Env is not able to mediate viral membrane fusion, it can participate in the transfer of viral particles by engaging CD4 in a virological synapse.

**Figure 8.**
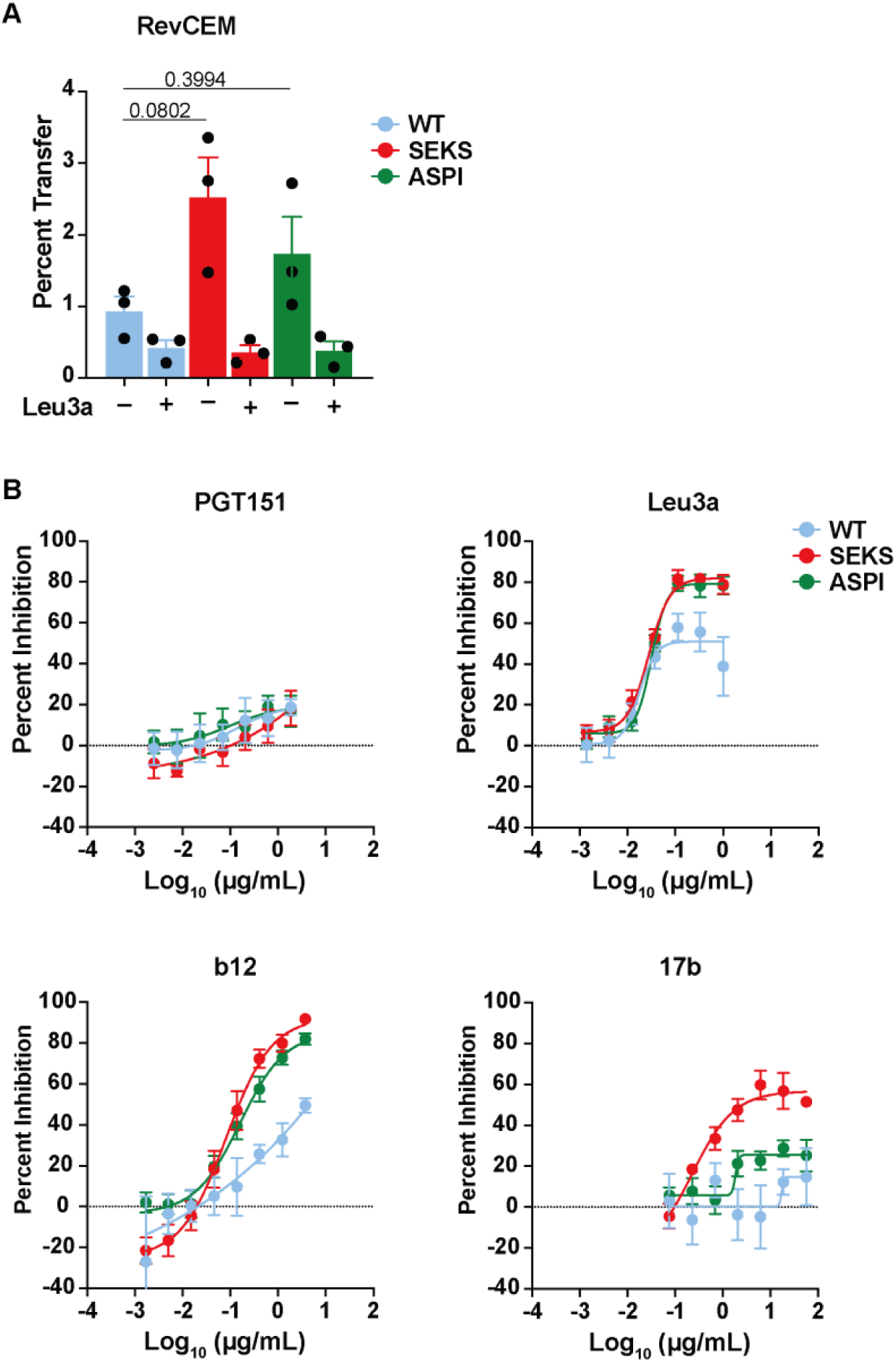
Uncleaved Env participates in CD4-dependent cell-to-cell transfer of HIV-1 to RevCEM. **(A)** Cell-to-cell transfer of WT and mutant GagiCherry_QH0692_ virions from nucleofected Jurkat T cells to RevCEM cells is shown. mCherry expression was used to normalize the nucleofection efficiency in donor cells before co-culturing for 6 hours, using flow cytometry. Leu3a at 1μg/mL was used as a negative control. p-values for Dunnett’s multiple comparisons test are shown. **(B)** Neutralization of WT and mutant NLCI_QH0692_ cell-to-cell transfer by PGT151, Leu3a, b12, and 17b. The donor cells were incubated with anti-Env mAbs and target cells with Leu3a for 30 minutes at 37°C before co-culturing cells. All experiments shown represent at least three independent experiments. The mean and SEM data are shown for each curve and bar graph.

Inhibition of cell-to-cell transfer was measured with a subset of the mAb panel previously discussed. PGT151 showed no activity against cell-to-cell transfer of the WT or mutant viruses (**Fig. 8B**). This is likely because PGT151 does not inhibit VS formation, as its epitope is not at the CD4 binding site. In contrast, b12, which recognizes the CD4 binding site, showed increased neutralization of SEKS- and ASPI-Env mediated cell-to-cell transfer (**Fig. 8B**). The increase in neutralization of cell-to-cell viral *transfer* by ASPI-Env relative to WT was similar in magnitude to that observed in cell-to-cell *infection*. We interpret this as indicating that the endocytic motif has a specific impact on the first step of cell-to-cell infection, i.e., VS formation and viral transfer. Similarly, 17b and Leu3a also showed enhanced neutralization of cell-to-cell transfer with SEKS-Env and ASPI-Env (**Fig. 8B**). These data further implicate uncleaved Env in participating in CD4-Env interactions that initiate the VS, despite not being fusogenic.

We also performed the cell-to-cell transfer assay with primary CD4_+_ T cells as target cells. Once more, incubation with Leu3a demonstrates a loss in transfer of fluorescent viral particles, highlighting the assays’ ability to measure VS-directed transfer of HIV-1 (**Fig. 9A**). The data shows that SEKS-Env can facilitate the transfer of viral particles to primary cells, at levels comparable to WT-Env (**Fig. 9A**). Additionally, ASPI-Env exhibited increased transfer compared to both WT- and SEKS-Env (**Fig. 9A**). We also observed that SEKS- and ASPI-Env increased sensitivity to b12 while no effect was seen with Leu3a (**Fig. 9B-C**). The sensitivity of cell-to-cell transfer, particularly to gp160-targeting antibodies, demonstrates that uncleaved Env can participate in the initial steps of cell-to-cell infection. Moreover, the similarity between SEKS- and ASPI-Env observed with Leu3a and b12 further supports our previous observations that the ratio of the cell surface Env has shifted to favor uncleaved when the main endocytic motif is mutated.

**Figure 9.**
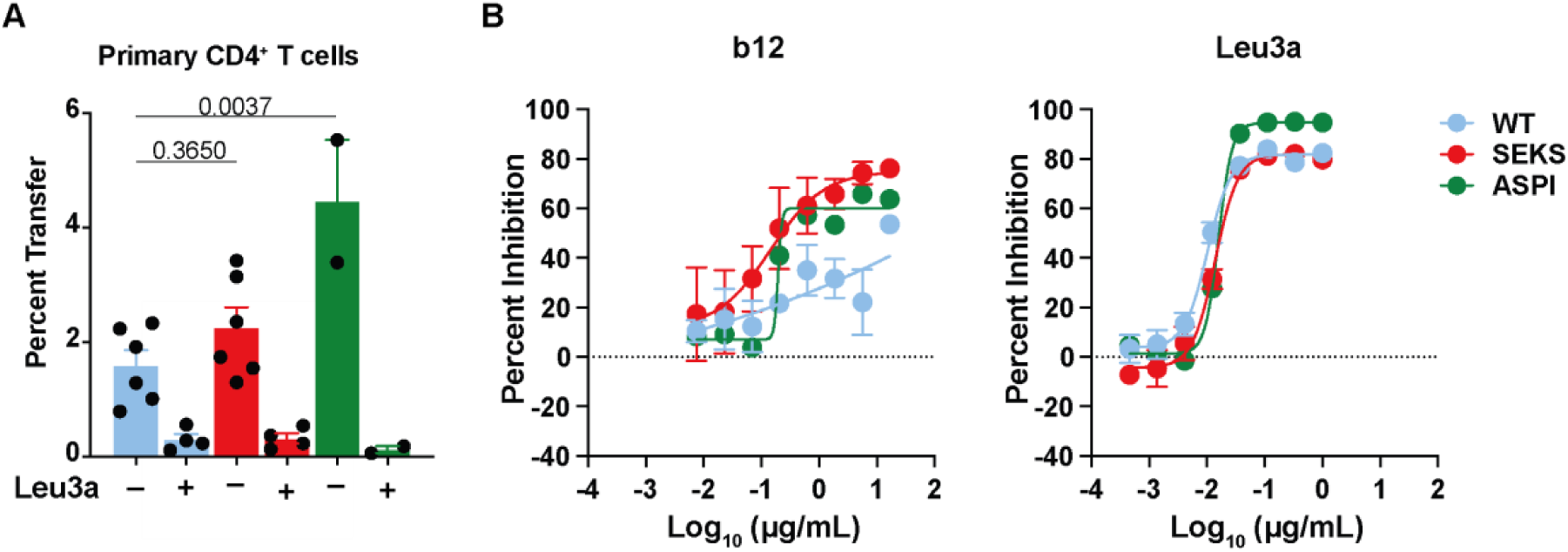
Uncleaved Env facilitates the cell-to-cell transfer of HIV-1 to primary CD4^+^ T cells. **(A)** Cell-to-cell transfer of WT and mutant GagiCherry_QH0692_ virions from nucleofected Jurkat T cells to primary CD4^+^ T cells is shown. mCherry expression was used to normalize the nucleofection efficiency in donor cells before co-culturing for 3 hours. Leu3a at 1μg/mL was used as a negative control. p-values for Dunnett’s multiple comparisons test are shown. **(B)** Neutralization of WT and mutant NLCI_QH0692_ cell-to-cell transfer by b12 and Leu3a. The donor cells were incubated with anti-Env mAb b12 and target cells with anti-CD4 mAb Leu3a for 30 minutes at 37°C before co-culturing cells. All experiments shown represent experiments performed with two different primary cell donors as target cells. The mean and SEM data are shown for each curve and bar graph.

## Discussion

A primary focus of vaccine development has been to induce responses that broadly recognize the native states of cleaved Env, which are present on mature virions. These strategies mainly aim to mimic the native Env present on virions, which is highly enriched for cleaved Env (29). There is comparatively less attention paid to the states of Env on the surface of infected cells, which can also participate in infection via cell-to-cell infection, yet which represents a mixture of uncleaved and cleaved Env (28, 29). Moreover, bNAbs, which are primarily characterized by cell-free infection assays, have demonstrated lower efficacy and potency against cell-to-cell infection compared to cell-free infection (3, 9–11). A potential mechanism proposed to explain the resistance to cell-to-cell infection is that the antigenically distinct forms of Env present on the cell surface, including both uncleaved and cleaved Env, compared to Env on the viral particle, contribute to HIV-1 infection and therefore require a more diverse Ab repertoire for inhibition (9, 13). We propose that uncleaved Env contributes to the increased antigenic diversity of Env on the cell surface and the increased resistance of cell-to-cell infection. Importantly, uncleaved Env has generally been overlooked as an active contributor to HIV-1 infection due to its inability to mediate fusion (62, 73) and has been proposed to primarily function as an immune decoy (16, 29, 77, 78). We hypothesized that the cleavage and glycosylation states of Env on the cell surface, as influenced by Env endocytosis, contribute to the sensitivity of cell-free and cell-to-cell infection to bNAbs. We found evidence that Env endocytosis can affect the abundance of cleaved Env on the cell surface with functional impacts on cell-free and cell-to-cell infection. Specifically, our data show that the ASPI mutation increases the sensitivity of cell-free infection to Abs that recognize uncleaved Env. The differences in cell-free infection may be more pronounced than those in cell-to-cell infection, in part because of the paucity of uncleaved Env on virions compared to the relative increase in uncleaved Env on the surface of infected cells. Because proteolytic processing affects glycosylation of Env during its synthesis, we also measured changes in glycosylation by measuring the sensitivity of specific lectins targeting oligomannose and complex glycans (22).

Endocytosed Env is thought to be recycled to particle assembly sites on the cell surface (44, 45). We did not observe a loss in the selective incorporation of cleaved Env onto virions with the endocytic mutant Env. Nevertheless, we do observe a slight increase in the packaging of an uncleaved Env glycoform (gp160h). Consistent with this, cell-free virus expressing ASPI-Env displayed enhanced sensitivity to antibodies that recognize uncleaved Env/more open conformations of Env. Additionally, 35O22, a mAb that is not cleavage-dependent but binds more efficiently when glycan processing is impaired, leading to an accumulation of high-mannose N-glycans (55), displayed enhanced binding to ASPI-Env. This further suggests that the endocytic mutant has also altered the glycan composition of Env packaged into virions, consistent with our lectin neutralization data.

While we acknowledge that uncleaved Env is not fusogenic and, therefore, cannot mediate cell-free infection, we propose that its increased incorporation into virions can still influence Ab neutralization and infectivity of HIV. Specifically, uncleaved Env may bind antibodies that sterically hinder cleaved Env from engaging with CD4 receptors, limit Env clustering required for fusion (79), and permit low-level cross-reactivity to cleaved Env, which can spontaneously sample conformations characteristic of uncleaved Env states (16). We note that the viral particles analyzed by western blotting in this study were derived from 293T cells. These cells are permissive to mutations in Env’s cytoplasmic tail (31, 80). Consequently, our findings may underestimate the true impact of the endocytic mutation on Env packaging, which in non-permissive cells may show a greater infectivity defect.

Endocytosed Env has been hypothesized to primarily be targeted for degradation through the endolysosomal pathway (81). Disrupting this pathway could therefore explain the alternative antigenic state of Env observed in our study. However, the fact that Env with a mutated endocytic motif can facilitate both cell-free and cell-to-cell infection does not suggest that ASPI-Env has led to an increase in misfolded or aggregated proteins.

Moreover, Hoffman et al. noted that while internalized Env colocalizes with endosome/lysosomal compartments, those compartments are not intended for degradation (82). Instead, they propose that these compartments are part of a T cell’s secretory pathway (82). Anand et al. also concluded that endocytosed Env-Ab complexes are not trafficked to the lysosome, having found no degradation of Env-Ab conjugates hours after internalization (78). Alternatively, uncleaved Env could also be targeted for degradation, considering its lack of fusogenic capability. Our data could support this possibility, as we observed a rise in the amount of uncleaved Env on the cell surface. However, no current data concludes that Ab-independent Env endocytosis is influenced by cleavage state. Additional studies are needed to probe further whether endocytosed Env is targeted for degradation, and whether Env’s cleavage state is a factor in the selective sorting to sites of viral assembly.

Endocytosed Env has also been proposed to traffic through the retrograde pathway to the trans-Golgi network (TGN) (46, 47). Considering the abundance of uncleaved Env present in the cell, trafficking to the Golgi could provide additional opportunities for cleavage, which is necessary for Env’s fusion potential (62, 73). Sequential trafficking through the Golgi would also allow for further glycosylation modifications to occur, which may not only contribute to building Env’s glycan shield but also influence the antigenic differences observed between Env on the virion and the cell surface. Alternatively, Zhang et al. demonstrated that glycans can help stabilize state 1 (83), which is favored by many bNAbs (14, 65, 84). Further studies are needed to determine whether internalized Env is trafficked to the TGN and whether Env transported through this pathway can be successively modified.

Our study also examined the effect of uncleaved Env on bNAb sensitivity in both cell-free and cell-to-cell infections. While uncleaved Env is not fusogenic and, therefore, incapable of mediating infection on its own (62, 73), the cell-to-cell transfer assay revealed that cleavage-deficient Env can facilitate the transfer of fluorescent viral particles. We attribute this to the SEKS-Env on a donor cell binding to a CD4 receptor on a target cell, initiating VS formation. The cell-free infection assay confirmed that SEKS-Env-derived viral particles are not infectious. However, viral particles have been found in intracellular compartments, suggesting that endocytic uptake of viral particles occurs, particularly at cell-to-cell contact sites (4, 8, 75, 76). Thus, the transfer assay measures the endocytic uptake of viral particles that takes place upon formation of the VS. Moreover, since Leu3a inhibited the transfer of SEKS-Env viral particles, we can further conclude that SEKS-Env mediates transfer by binding CD4 and triggering VS formation. This finding is supported by studies that have shown that cleavage-deficient Env can bind soluble CD4 (16, 74). Our neutralization data with b12 also suggest that the heterogeneous conformations of Env on the cell surface allow Env to engage in cell-to-cell infection. This conclusion is backed by existing literature showing that b12 exhibits a biphasic binding curve, indicating it recognizes binding sites with different affinities (85, 86). Since b12’s epitope overlaps with the CD4 binding site, b12 can inhibit viral fusion at the cell membrane and VS formation. Therefore, the nonsigmoidal neutralization curve could result from b12 targeting both uncleaved and cleaved Env, leading to different dissociation and binding rate constants. Additionally, while we observe an increase in uncleaved Env on the cell surface, and despite our panel having multiple antibodies that target epitopes more exposed on uncleaved Env, only b12 demonstrated enhanced potency against ASPI-Env. This further supports the need for an antibody that targets ENV-CD4 engagement to limit both virion attachment and VS formation.

A limitation of this study is that we focus on one primary isolate clone and test only a limited number of antibodies that target cleavage-sensitive epitopes. We note, however, that this study builds upon prior studies that investigated a larger number of epitopes, and we purposely focused on antibodies that are well-documented to bind cleaved or uncleaved Env preferentially. Additionally, the study of one T/F clone focuses on the question of whether Env endocytosis alters the cleavage ratio of cell surface Env. The main endocytic motif is well-conserved across HIV-1 lineages, suggesting its role in HIV-1 pathogenesis is universal (87). Nevertheless, additional studies must be conducted to confirm whether the effect of endocytosis on the conformation of cell surface and viral particle Env is broadly applicable to other T/F clones. An unexpected finding was the incomplete inhibition by Leu3a of cell-to-cell infection with WT Env. Li et al. previously demonstrated that Leu3a inhibits 100% of cell-to-cell infection (12); however, we only detected 80% inhibition at comparable concentrations. We note that these studies test different target cells, and there may be cell-type-dependent differences that allow CD4-independent entry of HIV-1.

In conclusion, we find that Env endocytosis is crucial for maintaining Env’s cleavage and glycosylation states during both cell-free and cell-to-cell infections, thereby contributing in some respects to the observed differences in sensitivities to bNAbs. This finding is significant because most vaccine strategies focus on inducing an immune response using antigens that mimic cleaved Env (88). Moreover, the current structure-based approaches employed to enhance Env expression remove or truncate the cytoplasmic tail (89–93). Based on the data presented here, we believe that limiting immunogens to cleaved Env may contribute to the ineffectiveness of current HIV-1 vaccines. Our study demonstrates not only that Env endocytosis is important to maintain the cleavage ratio of Env at the cell surface but also that uncleaved Env can participate in HIV-1 infection. Therefore, we propose that future vaccine studies and therapeutics consider targeting a broader range of conformations of Env that are represented on the cell surface and virions, and that they may include the endocytic recycling pathway to produce Env antigens.

## Acknowledgements

We are grateful to Dr. Markus Thali for the careful review of our manuscript, as well as to current and past members of the Chen Laboratory. We also thank the Flow Cytometry Core and Microscopy Core at the Icahn School of Medicine for their experimental guidance. This work was funded by the National Institutes of Health grant AI148064 to BKC.

